# Plant morphometry matches beetle diversity: effects of natural adjacent vegetation on grain Amaranth crop under small-holder conditions

**DOI:** 10.1101/2021.12.10.472071

**Authors:** Hugo Alejandro Álvarez, Gemma Clemente-Orta, Hortensia Carrillo-Ruiz, Jesús F. López-Olguín, Daniel Jiménez-García, Miguel A. Morón

## Abstract

Grain Amaranth, *Amaranthus hypochondriacus* L., is an emerging arable crop cultivated worldwide. One way to obtain resources from the crop, other than grain, is to grow Amaranths in the dry season and harvest only the leaves. In this environmental condition the response of an Amaranth agroecosystem to the presence of natural and semi-natural habitats or other crops has not been studied yet. We analysed the response of (1) Amaranth morphometry and (2) alpha and beta diversity of beetles to the nearness of adjacent vegetation and natural habitats (such as deciduous forest) at the small-holder conditions. Our results showed that *A. hypochondriacus* crop plants responded positively to the presence of an ecotone (adjacent vegetation) and the natural habitat (deciduous forest), i.e., *A. hypochondriacus* plants grew bigger in the section nearest to adjacent vegetation, which was a pattern consistent in time. Moreover, for beetles (considered as a bioindicator group), richness was different amongst the study areas and negatively followed the gradient of perturbation. These results suggest that Amaranth crop is sensitive to the presence of natural and semi-natural habitats but not to other crops in the dry season. In addition, beetles match the response of Amaranth plants. This is the first time that this type of data is recorded in grain Amaranth agroecosystem and it will help to understand the interaction amongst grain Amaranth agroecosystems, biodiversity, and natural adjacent vegetation to boost ecosystem services.

## Introduction

Grain Amaranth, *Amaranthus hypochondriacus* L., is an emerging arable crop, which is cultivated worldwide (Caselato-Sousa and Amaya-Farfán, 2012; Parra-Cota *et al.*, 2014). These herbaceous plants belong to the family Amaranthaceae in the Caryophyllales, and they are characterized by red to purple flowers in terminal or axial panicles (Alvarez *et al.*, 2017). The Amaranths are widely distributed in temperate zones (Mapes, 2001; Juan *et al.*, 2007; Mapes-Sánchez and Espitia-Rangel, 2010; Das, 2012). They are resistant to extreme climate and can grow in disturbed areas (Juan *et al.*, 2007; Mapes-Sánchez and Espitia-Rangel, 2010; Das, 2012).

It has been recorded that the harvest of Amaranth grain is an antique practice (e.g., Mexico, Aragón-García and Tapia-Rojas, 2009; Mapes-Sánchez and Espitia-Rangel, 2010; Perez-Torres *et al.*, 2011; Perez-Torres, 2012). An alternative way to obtain resources from this crop, other than grain, is to grow Amaranth in the dry season and harvest only the leaves, and thus, incorporating the leaves into meals (Hernandez and Herrerías, 1998; Pérez-Torres, 2012). Recently, the efficiency of grain Amaranth agroecosystems in terms of management and conservation has started to be studied, especially the implementation of organic technologies inside or next to the crop (Aragón-García and Tapia-Rojas, 2009; Alvarez *et al.*, 2016). However, under the extreme environmental conditions of dry season, the response of a grain Amaranth agroecosystem to the presence of natural adjacent vegetation and other crops has not been studied yet (but see Alvarez *et al.*, 2017, 2019c).

Agroecological theory suggests that natural adjacent vegetation reinforces microclimatic conditions in agroecosystems and provides food and shelter to wildlife that may increase the ecosystem functions (Altieri, 1999, 2000). However, not all organisms may respond in the same form to the presence of different types of vegetation, which it has been commonly related to the structure of the landscape and the amount of habitat fragmentation (Ries and Sisk, 2004; Laurance, 2007). Moreover, a community within an ecosystem under a transitional/colonization phase is different compared with a community in a well stablished ecosystem (Prach and Walker, 2011; Balmford *et al.*, 2012), this rationale can be extrapolated to agroecosystems separating annual and perennial crops. Recent accounts have shown that components of the surrounding agricultural matrix can be highly important to the ecology of agroecosystems because they can modulate the abundance of pests (i.e., arthropods or weeds) and their natural enemies (Tscharntke *et al.*, 2012, 2016; Sun *et al.*, 2015; Karp *et al.*, 2018; Clemente-Orta and Alvarez, 2019). For example, the presence of semi-natural habitats next or within orchards positively influence the abundance of pest predators (and sometimes the pests) (Wan *et al.*, 2019; Alvarez *et al.*, 2019a; 2019b). Conversely, due to intensive agricultural managements the presence of alfalfa crops nearby maize fields was more important than semi-natural habitats (Clemente-Orta *et al.*, 2020).

The aim of the present study was to assess the effects that (1) natural adjacent vegetation such as ecotones (semi-natural habitats) or conserved forest areas (natural habitats), and (2) other crops in adjacent growing areas may have on grain Amaranth crop in low-scale-farm (small-holder) conditions. We focused on the small-holder scale because it is the principal source of self-sustaining in most of the tropical and tempered zones worldwide. Thus, we analysed (1) plant morphometry in different zones within the crop at different developmental times, and (2) the structure of beetle alpha and beta diversity in different zones within the crop and adjacent vegetations, following a perturbation gradient. We hypothesised that natural adjacent vegetation will positively affect beetle diversity but not Amaranth growth.

## Material and methods

### Study area and experimental design

The study was carried out in the dry season of 2015 at San Francisco Huilango (14 Q 54.58.78-20.82.669), Puebla, Mexico. San Francisco Huilango is part of Tochimilco municipality that is at 1,860 m above sea level and has 800 to 1 300 mm of precipitation (INEGI, 2009; INAFED, 2010). The region has large surfaces of natural and conserved forest areas, which are near to the Popocatepetl volcano (Figure 1). For example, 43% of the region is occupied by temperate forest, 3% deciduous forest, and 3% grassland (INEGI, 2009; INAFED, 2010).

**Figure 1.**
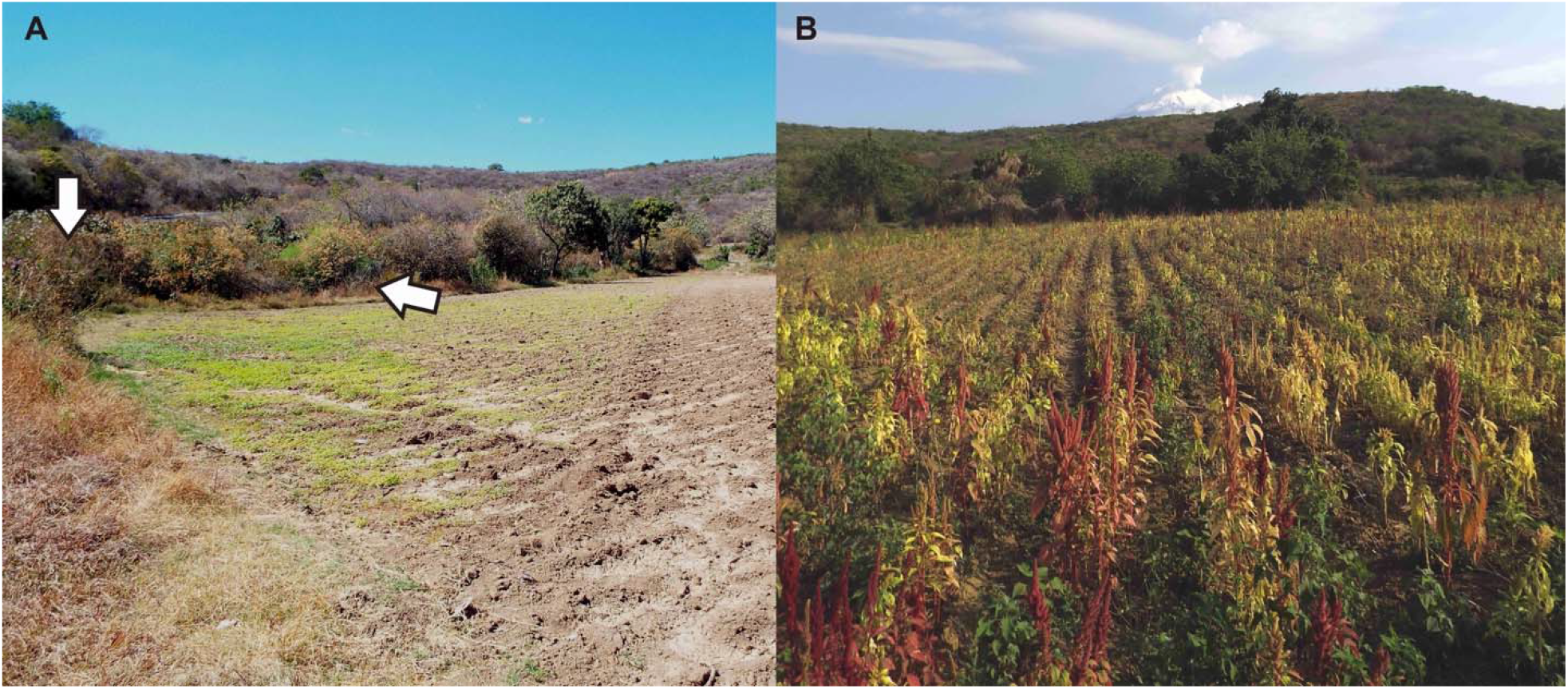
View of the study area near the Popocatepetl volcano. Natural adjacent vegetation as an ecotone next to the deciduous forest at the edge of the growing area (without a crop) (A). Arrows show the place of the ecotone. *Amaranthus hypochondriacus* crop at the end of the dry season (B).

We searched for a growing area located close to a natural system and an ecotone. We found a growing area that was approximately 2500 m^2^ and had at its west and southwestern edge an ecotone next to a deciduous forest (Figure 1). The rest of the perimeter adjoined with other growing areas and there was no presence of semi-natural vegetation between these areas.

An experiment with reduced management was developed. Firstly, the land was seeded with *A. hypochondriacus* in February 2015, then the land was fallow and plowed, and a manual weeding was done in April. We did not use any fertilizer or pesticide for pest control. Secondly, we delimited three plots of 300 m^2^ (20 × 15 m) each, with a separation of 100 m. Within each plot, we set three transects longitudinally (20 m) and each transect was placed 5 m apart. Finally, the plots were set (1) in the deciduous forest near the ecotone (natural area), (2) in the crop near the ecotone (hedge area), and (3) in the crop near the adjoining growing areas (non-hedge area).

### Plant morphometry

120 plants of *A. hypochondriacus* were randomly selected in the plots of the hedge area (*n* = 60) and non-hedge area (*n* = 60), i.e., we used only plants at the distal sections of each transect (*n* = 10) within plots. These sections were named section A (south section of the transect) and section B (north section of the transect). Four morphometric variables, stalk width, leaf width, panicle length, and plant height were measured linearly for each plant with (1) vernier callipers (accuracy of 0.01 mm) and (2) a measuring tape (accuracy of 0.1 cm). Stalk width was measured transversally at the middle point of the stalk (steam). Leaf width was measured transversally from edge to edge at the widest point of the leaf. Panicle length was measured from the base of the panicle to the distal point. Plant height was measured from the base of the stalk to the apical point of the panicle. We made the measurements in two times: after two months of sowing (time 1) and after four months of sowing (time 2).

### Beetle diversity

We sampled from April to July (once a month) the foliage of Amaranth crop plants placed in the transects within the plots of the three study areas following the methodology of Morón and Terrón (1988). For example, beetles of the foliage were collected by batting plants with an entomological net in five randomly distributed battings (one sample) per area. Samples were collected from 900 to 1300 hours every 20 days. Specimens were preserved in 70% alcohol and transported to the laboratory for identification. All specimens were identified to species level (otherwise specified) and deposited in the entomological collection of the Institute of Research in Natural Sciences and Humanities (IINCINH A.C.).

### Data analysis

Data of morphometric measurements of *A. hypochondriacus* showed no normal distribution (Shapiro-Wilk normality test: Stalk W = 0.877, *p* = 0.001; leaf W = 0.942, *p* = 0.001; panicle W = 0.829, *p* = 0.001; height W = 0.909, *p* = 0.001). Thus, the correlation structure amongst all morphometric variables was examined by calculating the pairwise Pearson’s product-moment correlation coefficients. Then, we fitted generalized linear models (GLM) for each of the four variables. We used each measurement variable as the dependent variable and time, area, and section as factors. Interactions amongst factors were introduced in the analysis.

Data of beetle richness and abundance were used to assess alpha and beta diversity amongst the study areas. First, to analyse alpha diversity we calculated the (1) Shannon and (2) Simpson diversity indexes for each area. Then, the species accumulation curves and non-parametric estimators were calculated for the three areas together. Finally, the sampling effort was calculated for each area. Alpha diversity analyses were computed using the software EstimateS, v 9.1.0 (Colwell, 2013) and SPADE (February 2009 actualization, Chao and Shen, 2009). Second, we analysed beta diversity searching for the replacement in the form of richness gradients. We used the index of multiple-site dissimilarities *(Msim)* of Baselga *et al.* (2007), which is calculated:

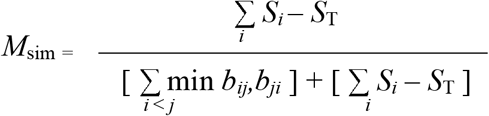

where *S_i_* is the richness in site *i* (α), *S*_T_ is the richness of the total of sites (*γ*), *b_ij_* indicates the number of species that only occurred in site *i* but not in site *j, b_ji_* indicates the number of species that only occurred in site *j* but not in site *i*, and min is the minimum number between *b_i_j* and *b_ij_*.

The index of multiple-site dissimilarities is of much help and confidence, because species replacement/turnover and species loss could be the cause for the difference in species composition amongst sites in a richness gradient. The index only responds to species replacement and does not depend on differences of species richness amongst sites, which makes it have an advantage over other indexes (Baselga, 2007; Baselga *et al.*, 2007; 2010; 2012). We used the function “beta.multi” from package “betapart” (Baselga *et al.*, 2015) to compute the index of multiple-site dissimilarities. This function drops a list of values of similarity: (1) the value of the components of the replacement, measured as Simpson’s similarity; (2) the value of the nesting components, measured as the nesting fraction resulting from Sorensen’s similarity; and (3) the value of overall beta diversity, measured as Sorensen’s similarity. The possible values go from zero to one, where one indicates the maximum possible value (Baselga *et al.*, 2015).

The software, R v. 3.3.1 (R Developmental Core Team, 2015) was used to compute correlation analysis and to compute the index of multiple-site dissimilarities. The software SPSS, v.19 (SPSS Inc.) was used to compute GLMs.

## Results

### Plant morphometry

Correlation analysis showed that all variables are positively correlated. The variable that had the greatest correlation coefficients with all the variables was height (Table 1). Height and panicle showed the greatest positive correlation coefficients, with the exception of hedge area in time one (Table 1). Conversely, stalk showed the lowest correlation coefficients.

**Table 1.**
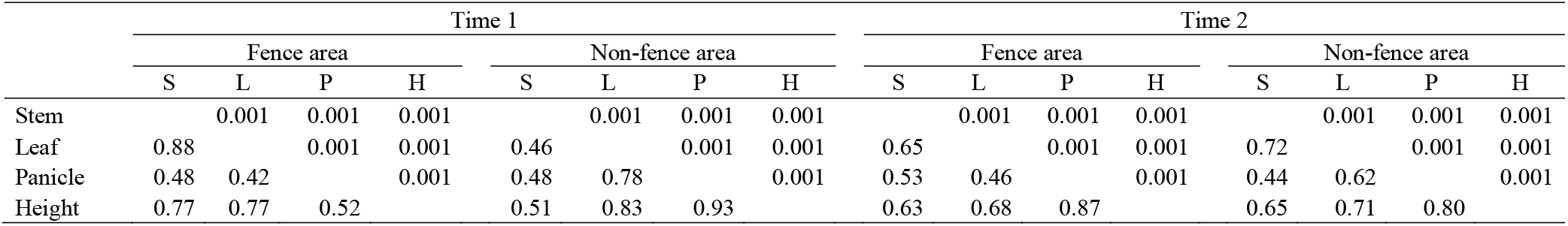
Statistics of pairwise Pearson’s product moment correlation. It is shown below the diagonal correlation coefficients and above the diagonal *p* values of each morphometric variable according to time and area.

|  | Time 1 |  |  |  |  |  |  |  | Time 2 |  |  |  |  |  |  |  |
| --- | --- | --- | --- | --- | --- | --- | --- | --- | --- | --- | --- | --- | --- | --- | --- | --- |
|  | Fence area |  |  |  | Non-fence area |  |  |  | Fence area |  |  |  | Non-fence area |  |  |  |
|  | S | L | P | H | S | L | P | H | S | L | P | H | S | L | P | H |
| Stem |  | 0.001 | 0.001 | 0.001 |  | 0.001 | 0.001 | 0.001 |  | 0.001 | 0.001 | 0.001 |  | 0.001 | 0.001 | 0.001 |
| Leaf | 0.88 |  | 0.001 | 0.001 | 0.46 |  | 0.001 | 0.001 | 0.65 |  | 0.001 | 0.001 | 0.72 |  | 0.001 | 0.001 |
| Panicle | 0.48 | 0.42 |  | 0.001 | 0.48 | 0.78 |  | 0.001 | 0.53 | 0.46 |  | 0.001 | 0.44 | 0.62 |  | 0.001 |
| Height | 0.77 | 0.77 | 0.52 |  | 0.51 | 0.83 | 0.93 |  | 0.63 | 0.68 | 0.87 |  | 0.65 | 0.71 | 0.80 |  |

Table 2 summarizes the statistics of each morphometric variable in base to experimental design and GLMs. According to the analyses, variables showed differences within each factor. Variables were bigger at time two, except for leaf that showed a decrease in length. Hedge area showed greater sizes than the non-hedge area. In addition, section A showed larger sizes than section B. There were interactions within all the variables, i.e., between factors time and type, and between factors type and section. The interaction time-section was significant for the variable panicle. This indicates that panicle was big in section A, but it was bigger at time two, i.e., it grew exponentially and duplicated its size (Table 2). The interaction type-section showed that section A in the hedge area had bigger sizes. Conversely, the rest of the sections were similar amongst each other with low sizes. Overall, the patterns were stronger in panicle and height (Figure 2).

**Figure 2.**
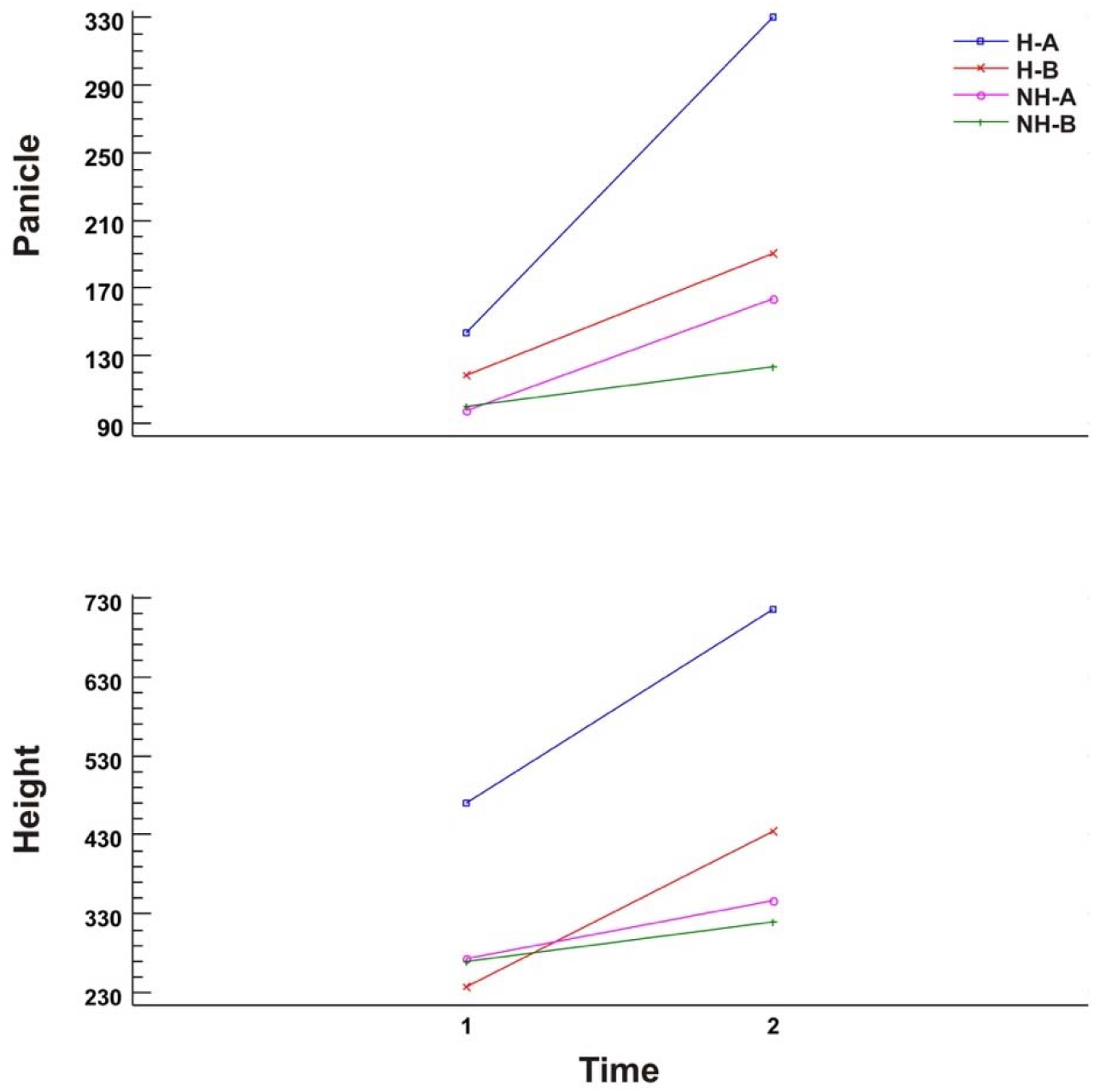
Differences in growth of *Amaranthus hypochondriacus* traits. Panicle and height mean (centimetres) in each section per area (area-section), in the two sample periods. Hedge area (H), Non-hedge area (NH).

**Table 2.** Statistics of *Amaranthus hypochondriacus.* Descriptive statistics: mean and standard deviation of morphometric variables. Model F statistics and *p* values according to each GLM results. The first group of data shows comparison of treatments into each of the factors: time, area, and section. The second group of data shows interactions amongst factors.

|  | STALK |  |  |  |  | LEAF |  |  |  | PANICLE |  |  |  | HEIGHT |  |  |  |
| --- | --- | --- | --- | --- | --- | --- | --- | --- | --- | --- | --- | --- | --- | --- | --- | --- | --- |
|  | *n* | Mean | SD | *F* | *p* | Mean | SD | *F* | *p* | Mean | SD | *F* | *p* | Mean | SD | *F* | *p* |
| TIME |  |  |  |  |  |  |  |  |  |  |  |  |  |  |  |  |  |
| 1 | 120 | 4.34 | 1.70 | 21.89 | 0.001 | 28.11 | 9.73 | 5.46 | 0.019 | 114.85 | 121.01 | 32.82 | 0.001 | 312.36 | 186.20 | 35.84 | 0.001 |
| 2 | 120 | 5.51 | 2.39 |  |  | 25.18 | 12.18 |  |  | 201.70 | 124.01 |  |  | 454.00 | 224.82 |  |  |
| TYPE |  |  |  |  |  |  |  |  |  |  |  |  |  |  |  |  |  |
| Fence | 120 | 5.65 | 2.32 | 33.05 | 0.001 | 31.08 | 11.86 | 46.15 | 0.001 | 195.47 | 156.46 | 24.50 | 0.001 | 464.04 | 257.53 | 45.74 | 0.001 |
| No Fence | 120 | 4.20 | 1.69 |  |  | 22.21 | 8.18 |  |  | 121.08 | 80.97 |  |  | 302.33 | 125.72 |  |  |
| SECTION |  |  |  |  |  |  |  |  |  |  |  |  |  |  |  |  |  |
| A | 120 | 5.34 | 2.37 | 11.54 | 0.001 | 29.49 | 12.68 | 20.12 | 0.001 | 183.52 | 136.48 | 11.59 | 0.001 | 450.87 | 250.20 | 32.94 | 0.001 |
| B | 120 | 4.50 | 1.82 |  |  | 23.80 | 8.38 |  |  | 133.04 | 117.96 |  |  | 315.50 | 153.25 |  |  |
| TIME/TYPE |  |  |  |  |  |  |  |  |  |  |  |  |  |  |  |  |  |
| 1/Fence | 60 | 4.80 | 1.75 | 5.04 | 0.025 | 30.67 | 11.25 | 10.06 | 0.002 | 130.91 | 156.77 | 8.93 | 0.003 | 353.21 | 231.33 | 14.46 | 0.001 |
| 1/No Fence | 60 | 3.88 | 1.53 |  |  | 25.54 | 7.13 |  |  | 98.80 | 66.58 |  |  | 271.51 | 114.10 |  |  |
| 2/Fence | 60 | 6.50 | 2.51 |  |  | 31.49 | 12.53 |  |  | 260.03 | 127.74 |  |  | 574.86 | 235.11 |  |  |
| 2/No Fence | 60 | 4.52 | 1.79 |  |  | 18.88 | 7.84 |  |  | 143.37 | 88.22 |  |  | 333.15 | 130.23 |  |  |
| TIME/SECTION |  |  |  |  |  |  |  |  |  |  |  |  |  |  |  |  |  |
| 1/A | 60 | 4.84 | 2.00 | 0.51 | 0.472 | 30.67 | 11.21 | 0.23 | 0.628 | 120.36 | 108.66 | 7.8 | 0.005 | 370.91 | 218.15 | 0.77 | 0.378 |
| 1/B | 60 | 3.84 | 1.15 |  |  | 25.54 | 7.20 |  |  | 109.35 | 132.91 |  |  | 253.81 | 123.96 |  |  |
| 2/A | 60 | 5.84 | 2.61 |  |  | 28.31 | 13.99 |  |  | 246.67 | 132.86 |  |  | 530.83 | 256.27 |  |  |
| 2/B | 60 | 5.17 | 2.10 |  |  | 22.05 | 9.15 |  |  | 156.73 | 96.20 |  |  | 377.18 | 155.77 |  |  |
| TYPE/SECTION |  |  |  |  |  |  |  |  |  |  |  |  |  |  |  |  |  |
| Fence/A | 60 | 6.54 | 2.38 | 15.75 | 0.001 | 36.68 | 12.22 | 21.27 | 0.001 | 236.55 | 150.56 | 5.05 | 0.025 | 592.43 | 255.20 | 32.08 | 0.001 |
| Fence/B | 60 | 4.76 | 1.88 |  |  | 25.48 | 8.42 |  |  | 154.40 | 152.56 |  |  | 335.65 | 187.40 |  |  |
| No Fence/A | 60 | 4.15 | 1.66 |  |  | 22.31 | 8.35 |  |  | 130.49 | 95.83 |  |  | 309.31 | 142.77 |  |  |
| No Fence/B | 60 | 4.25 | 1.73 |  |  | 22.11 | 8.07 |  |  | 111.68 | 62.14 |  |  | 295.35 | 106.89 |  |  |

### Beetle diversity

We collected 21 beetle species of the families Brentidae, Chrysomelidae, Coccinelidae, Melyridae, Phengodidae and Scarabaeidae. Table 3 shows the relative abundance of beetle species, taxonomic information, and occurrence in the different study areas. The family Chrysomelidae had the biggest richness with 12 species, followed by the family Brentidae with three species. Families Coccinelidae and Melyridae were represented by two species and the families Phengodidae and Scarabaeidae by one species. *Diabrotica balteata* LeConte, was the most abundant species followed by *Zygogramma signatipennis* (Stål), *Disonycha melanocephala* Jacoby, and *Coelocephalapion buchanani* (Kissinger).

**Table 3.**
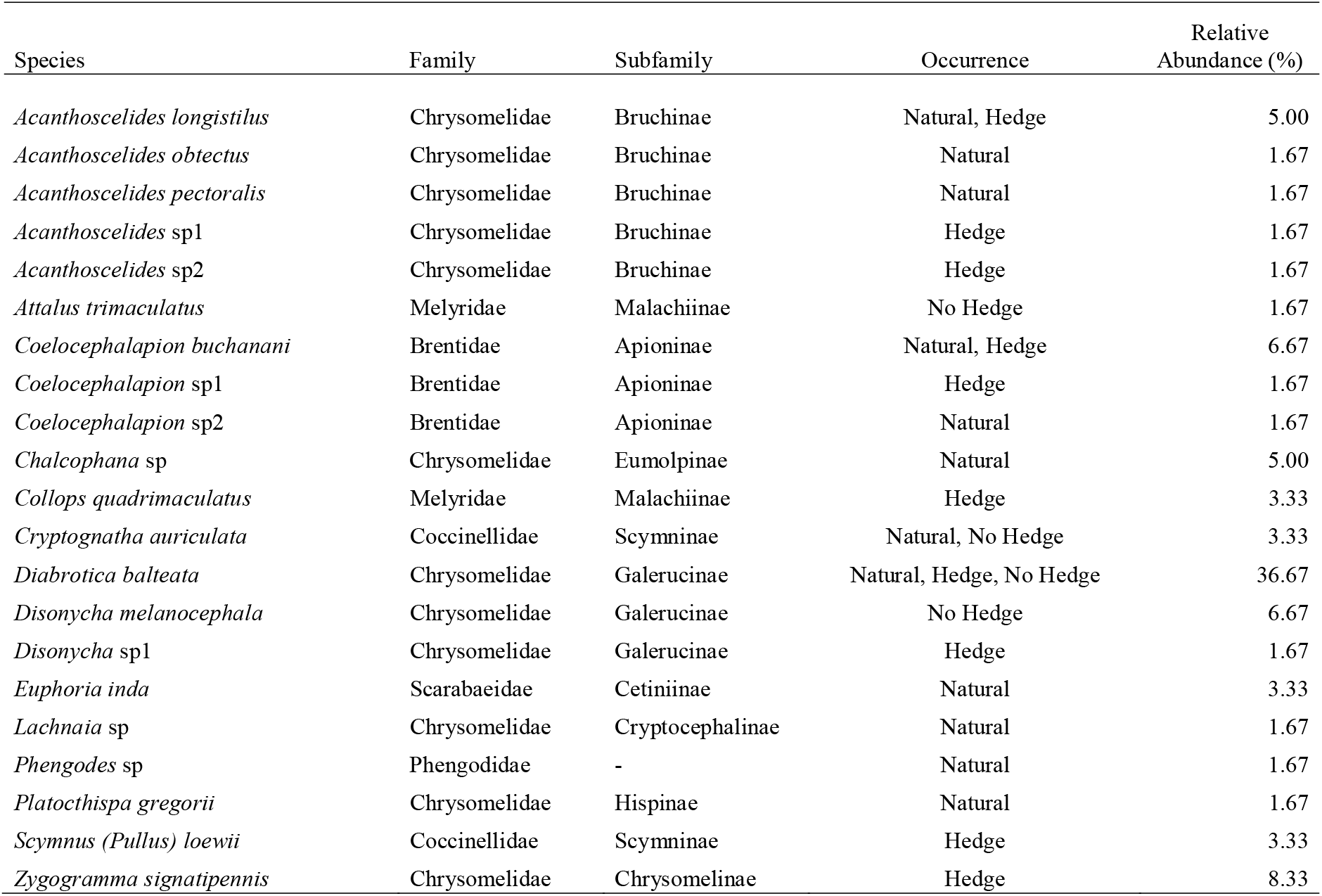
Relative abundance, taxonomic information, and occurrence of beetle species associated with *Amaranthus hypochondriacus* and deciduous forest. Study areas: Natural area, hedge area, and non-hedge area.

Figure 3 shows the calculated species accumulation curve for the overall data with 95% upper and lower values. The non-parametric estimators: ACE, Chao 1, Chao 2, Jackknife 1, Jackknife 2, and ICE were visualized using the data obtained in the software EstimateS. The non-parametric estimators that better adjusted to the asymptote were ACE and Chao 1 (Figure 3). These two non-parametric estimators are based on abundance. Specifically, Chao 1 is based on the number of rare species (singletons and doubletons) but in ACE the rare species are defined by the maximum number in abundance.

**Figure 3.**
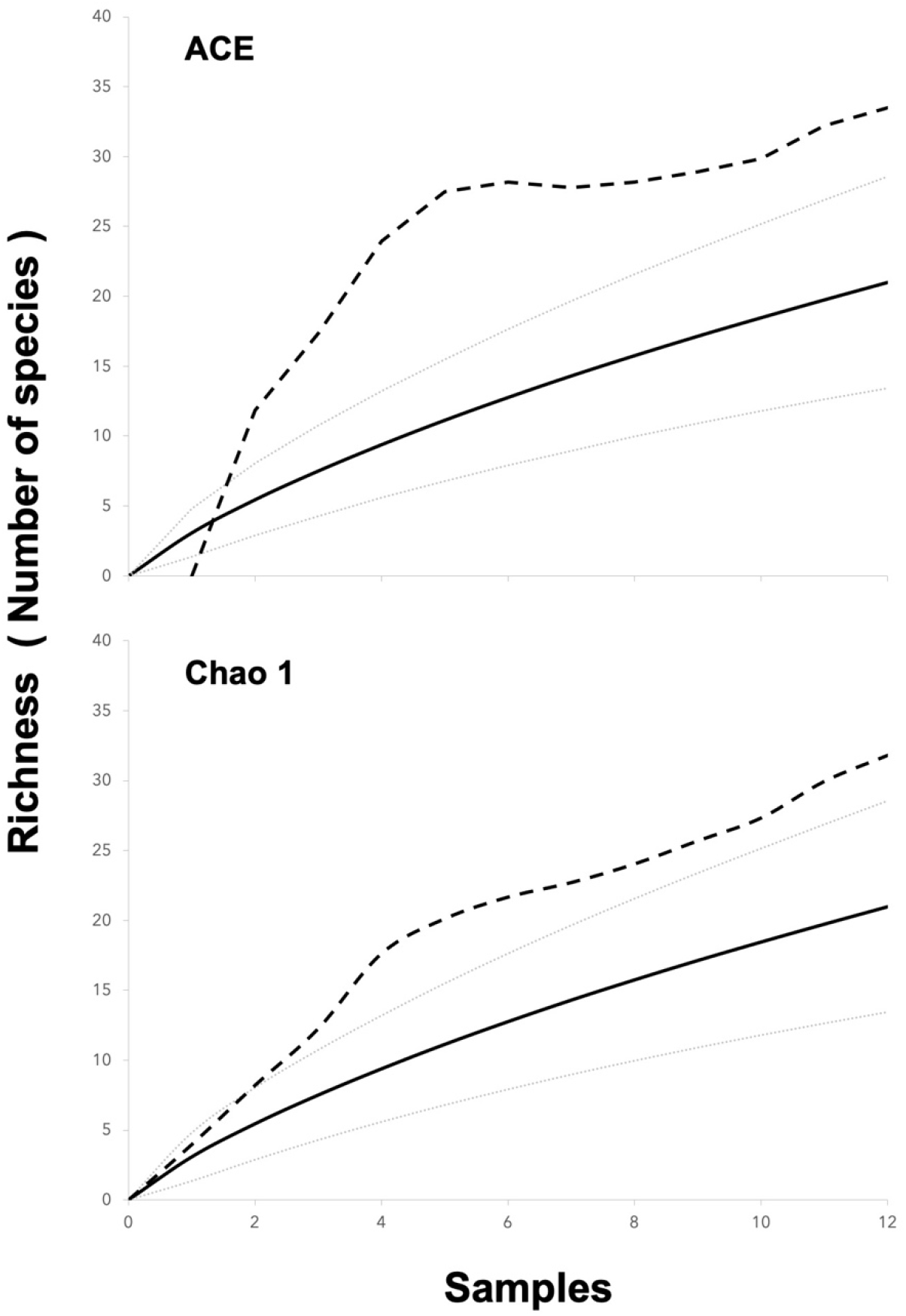
Calculated species accumulation curve of beetle species richness (solid lines) with 95% upper and lower CI for both non-parametric estimators (dotted lines): ACE and Chao1.

The calculated sampling effort, based on Chao 1 and ACE, was 80% for hedge area, 80% for non-hedge area, and 60% for the natural area. Moreover, the diversity indexes of Shannon (H’) and Simpson (D) were high for the natural area (H’ = 2.370; D = 0.10494) followed by the hedge area (H’ = 2.013; D = 0.17202). Conversely, the indexes were low for the non-hedge area (H’ = 0.955; D = 0.48097).

The index of multiple-site dissimilarities showed that there was high similarity amongst sites (components of the replacement value = 0.705), however, there was no nestedness amongst sites (nesting components value = 0.094). The analysis showed that the beta diversity of the community was high (overall beta diversity value = 0.8) (Figure 4).

**Figure 4.**
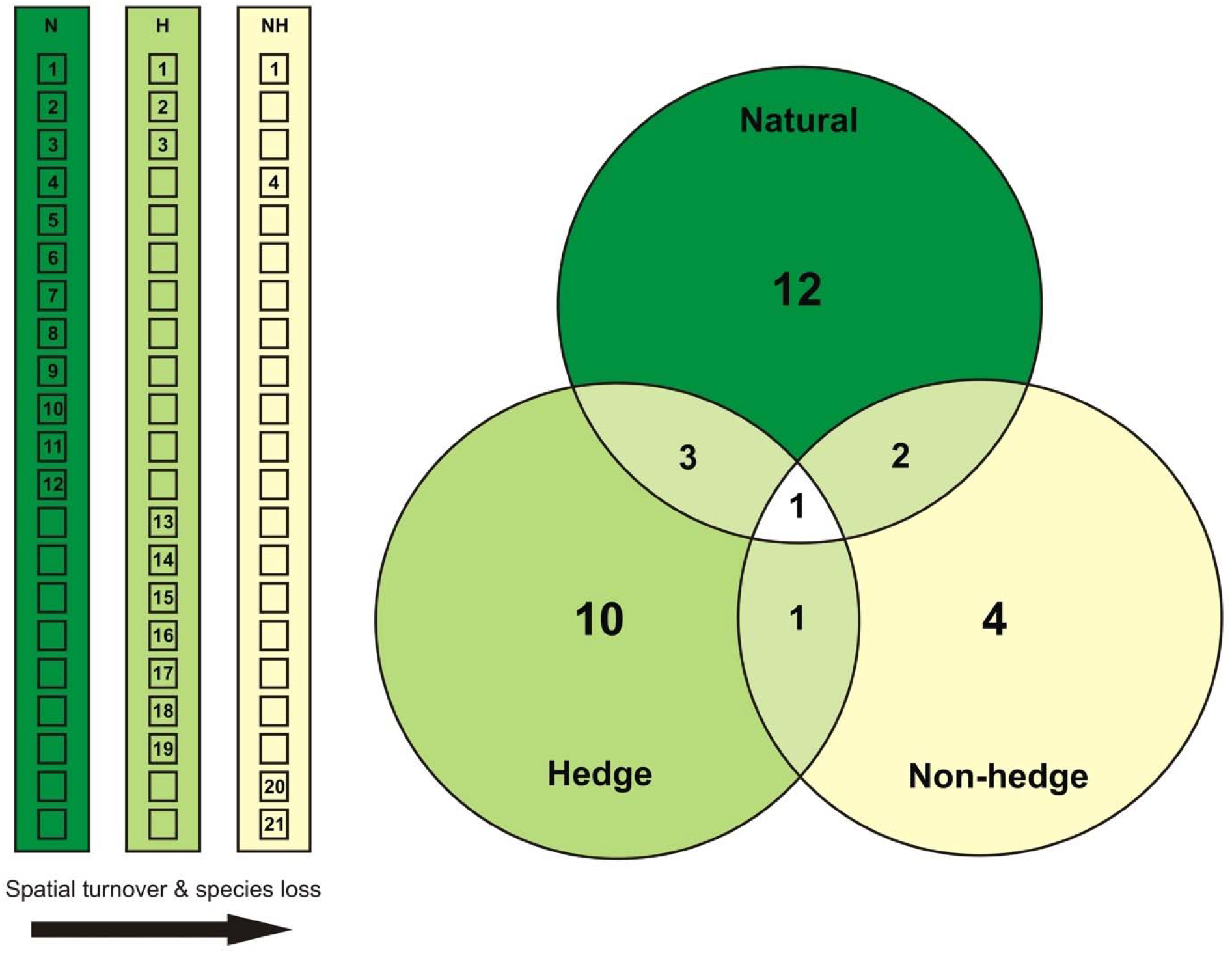
Graphic representation of the nestedness and turnover of beetle community associated with *Amaranthus hypochondriacus* and the deciduous forest. Natural system (N), hedge area (H), and non-hedge area (NH). Rectangles represent the ecological gradient amongst study areas. Numbers inside squares represent beetle species arranged from most to least common and separated by abundance. Circles represent the ecological community, showing the number of species in each of the study areas and species shared between areas.

## Discussion

Edge effects have been defined as changes in biotic or physical variables occurring at the transition between adjoining habitats (Murcia, 1995; Lopez-Barrera *et al.*, 2007). Our results showed that beetles (considered as a bioindicator group) and *A. hypochondriacus* crop plants responded positively to the presence of an ecotone, which can be interpreted as positive edge effects.

According to the development of our experiment, the pattern of growth of the traits of *A. hypochondriacus* plants was similar in hedge and non-hedge areas. However, traits within the section A of the hedge area, which is the nearest point to the ecotone, grew bigger. This pattern was consistent in time, but panicle and plant height showed the biggest effects (Figure 2). It is known that plant growth depends on the conditions in a growing area (Hopkins and Hüner, 1995), and recent accounts suggest that in some scenarios high-diversity plant assemblages benefit native plant-invaders rather than alien invaders, which is due to the biogeographical history of plants species (Sun *et al.*, 2015). If we accept that native plant species have better opportunities to settle when they are invading a new territory, such as an agriculture matrix, then these species will be beneficiated for being near to a native ecotone or hedge. This could be the case of *A. hypochondriacus,* because this species of Amaranth is a native species in Mesoamerican zones and it has been cultivated in the study region for centuries (Mapes-Sánchez and Espitia-Rangel, 2010).

On the other hand, beetle diversity negatively followed the perturbation gradient, i.e., a gradient related with the quality of the habitat (ecological habitats, Colwell *et al.*, 2004; Desrochers and Anand, 2005). Indeed, the natural area (deciduous forest) was the most diverse followed by the hedge area and finally the non-hedge area. Furthermore, species richness had a transitional response, i.e., richness responded to the presence of natural adjacent vegetation in a negative form within the natural system (high quality habitat) and in a positive form within the crop (low quality habitat). Interestingly, the species were not nested, therefore, the communities with low richness are not subsets of the communities with high richness (Figure 4). Only *A. longistilus, C. buchanani, C. auriculata,* and *D. balteata,* were present in the three areas.

Nonetheless, our results showed that the calculated sampling efficiency in the natural system is low, 12 species were collected in this area. This could be the result of the presence of several singletons that were collected in the final period of the sampling season, which was the beginning of the rainy season. Beetle adults in this area of the country usually emerge in May or June (Lobo and Morón, 1994; Pérez-Torres *et al.*, 2011; M.A. Morón personal comments). It is possible that the adults collected at the end of the sampling season belong to the beetle fauna of the rainy season.

Overall, our three study areas form a matrix of highly different properties. According to the rationale in Duelli *et al.* (1990), the growing area in this study had the properties of a hard edge but the ecotone (adjacent vegetation) had the properties of a soft edge. Thus, we can explain the positive edge effects seen in plants growth and beetle diversity in the context of a resource competition framework (Sun *et al.*, 2015) and/or a resource distribution model (Ries and Sisk, 2004; Macreadie *et al.*, 2010). For example, complementary resource distribution refers to a scenario where two different adjacent habitats have different resource availability (quantity and quality). When these habitats are highly different a complementary distribution of resources will drive a positive response, i.e., the low-quality habitat will have lesser abundance of animals than the other habitats but the individuals at the boundaries will benefit due to the availability of new resources. The resource-based model shows that resources could be concentrated at an edge, therefore when there is fragmentation of natural habitats resources are better exploited around edges by native and/or opportunistic species (Ries and Sisk, 2004; Macreadie *et al.*, 2010), which could be the case of *A. hypochondriacus* crop.

This study is a first step to understand the interaction amongst grain Amaranth agroecosystems, biodiversity, and natural or semi-natural habitats. The prescence of a specific pattern at local scale, suggest that a landscape-scale perspective is needed in forthcoming research to prove if the response to the proximity to natural adjacent vegetation, showed here by *A. hypochondriacus,* is a rule in grain Amaranth agroecosystems.

## Disclosure

The authors are not aware of any affiliations, memberships, funding, or financial holdings that might be perceived as constituting a conflict of interest.

## Notes

### Competing Interest Statement

The authors have declared no competing interest.

